# Expected differences in diversity and rarity between communities containing sexually versus asexually reproducing taxa

**DOI:** 10.1101/397174

**Authors:** Cristina M. Herren

## Abstract

Recent studies evaluating the community structures of microorganisms and macro-organisms have found greater diversity and rarity within micro-scale communities, compared to macro-scale communities. However, reproductive method has been a confounding factor in these comparisons; the microbes considered generally reproduce asexually, while the macro-organisms considered generally reproduce sexually. Sexual reproduction imposes the constraint of mate finding, which can have significant demographic consequences by depressing birth rates at low population sizes. Here, I examine theoretically how the effects of mate finding in sexual populations translate to the emergent community properties of diversity, rarity, and dominance. Using continuous-time Markov chain models, I compare communities with and without constraints of mate limitation. In mate-limited sexual populations, the decreased growth rates at low population densities translates to a much higher extinction rate. In communities consisting of sexually reproducing taxa, the increased extinction rate due to mate limitation decreases expected diversity. Furthermore, mate limitation has a disproportionately strong effect on taxa with low population density, leading to fewer rare taxa. These shifts in community structure mirror recent empirical studies of micro versus macro communities, indicating that reproductive method may contribute to observed differences in emergent properties between communities at these two scales.

## Introduction

Ecologists have historically been fascinated by the diversity of microbial communities (Hutchinson 1961), and several recent studies have indeed demonstrated differences in community structure between microbes and larger “macro” organisms (Nemergut et al. 2013, Hansen and Carey 2015, Locey and Lennon 2016, Shoemaker et al. 2017, Meyer et al. 2018). Generally, microbial communities have higher diversity that results in part from the large number of rare taxa (Neufeld and Lynch 2015). But, other properties, such as abundance of the most dominant taxon, are indistinguishable between communities at the two different scales (Locey and Lennon 2016). Despite increasing data on which to base these comparisons, the mechanisms generating these patterns of population distributions within and between communities are poorly understood (Shade 2017). One prominent additional difference between many of the microbial populations and macro populations in prior comparative studies is reproductive method; the microbial populations considered (bacteria, archaea, and most phytoplankton) reproduce asexually, while most macro populations considered have sexual reproduction. Here, I examine theoretically whether reproductive method can contribute to observed differences in community structure between asexually reproducing microorganisms and sexually reproducing macro organisms.

Individuals in sexually reproducing populations must encounter a mate before reproducing, whereas asexual individuals do not have this constraint. Mate finding and its consequences on population dynamics have been extensively studied in the theoretical literature (beginning with Volterra 1938), in part because it is one mechanism that causes Allee effects (reviewed in Gascoigne et al. 2009). An Allee effect is defined as positive density dependence within a population, meaning that individual-level growth rates increase as population density increases (Odum and Allee 1954). When an Allee effect is present, the benefit of encountering another individual from the population outweighs negative interactions, such as competition, and individuals become more reproductively successful as density increases (Courchamp et al. 1999). In populations with sexual reproduction, sparse populations are slow growing due to the inability to find a mate. Mate encounters become more frequent as the population grows, such that per-capita fitness increases as density increases. The effects of mate finding on population growth are prominent when population sizes are small, but decrease when population size is large and mates are no longer limiting (Dennis 1989).

Many previous theoretical models have considered mate finding and Allee effects using differential or difference equations describing the population growth rate (Odum and Allee 1954, Dennis 1989, Boukal and Berec 2002). Strong reductions in birth rates due to mate limitation can cause population declines at low abundance, effectively setting a “critical density” below which the population becomes extinct (Gerritsen 1980). When the population size is greater than the critical density, the population continues to grow until reaching a stable equilibrium at its carrying capacity (Stephens et al. 1999). However, a major drawback of deterministic models is the inability to consider time to extinction for populations with a positive stable equilibrium; with deterministic equations, any population with a positive stable state will persist indefinitely. This result conflicts with the empirical observation that smaller populations are more vulnerable to extinction (Purvis et al. 2000).

Stochastic models are promising for studying demographic consequences of mate finding, because they allow for extinction in populations that would otherwise reach a positive carrying capacity (Lande 1993). Whereas the persistence of populations in deterministic equations is governed by local population growth rates around an equilibrium, persistence in stochastic models depends on the growth rates at every density (Assaf and Meerson 2010). In other words, the chance of extinction in real populations is related to population growth rates near zero, which are important in stochastic models but rarely considered in deterministic models. Several forms of stochastic populations models have been used to study populations with Allee effects, often with discrete time models (Stephan and Wissel 1994, Allen et al. 2005, Sun 2016). These studies have concluded that diminished growth rates at low population densities can substantially decrease expected time to extinction (Stephan and Wissel 1994, Dennis 2002). However, it is computationally difficult to model multiple interdependent populations or populations with overlapping generations in discrete time models (Allen and Allen 2003), which is often a prohibitive barrier to such studies.

Here, I compare population and community dynamics between communities that must find mates before reproducing and communities where populations have no mate limitation (a case equivalent to asexual reproduction). I use stochastic models to evaluate demographic consequences of mate finding. First, I use continuous-time Markov chain models (CTMCs) to study how mate limitation alters time to extinction for single populations. These models use a computationally efficient simulation algorithm, which allows for simulation of multiple coexisting populations. Such models have been extensively used for simulating chemical reaction networks (Gillespie 2007), but can also be used for modeling population dynamics (Dobramysl et al. 2018). After obtaining the mean times to extinction obtained from these models, I use the island biogeography framework to evaluate how varying extinction times translate to changes in community diversity. The island biogeography framework posits that the expected long-term community diversity can be calculated by identifying the number of taxa where immigration and extinction rates are equal (MacArthur and Wilson 1967). In these models, I assume identical immigration rates between the various communities, but extinction rates are a function of mate limitation. Finally, I simulate communities consisting of populations with differing growth rates to evaluate how consequences of mate limitation scale to heterogeneous communities. I show that the constraint of mate search decreases diversity, primarily by excluding rare taxa, whereas dominance of the most abundant population is unaffected.

## Methods

### Single population dynamics

First, I studied the effects of mate limitation on the time to extinction for single populations. I used CTMCs to evaluate time to extinction, implemented with the Gillespie algorithm (Gillespie 1977). Briefly, these models record births and deaths in a population as events that occur with varying frequency, depending on population size. Births are marked by the addition of a single individual to the population, whereas deaths remove a single individual. The overall rate at which any event (birth or death) occurs is the sum of the birth and death rates. The time until the next event is exponentially distributed with a parameter equal to the summed event rates. Therefore, as event rates increase, waiting time until the next event decreases. After drawing a random value from the exponential distribution for the time increment, the magnitudes of the instantaneous birth and death rates indicate whether a birth (add one individual) or death (remove one individual) is more likely to occur. Another random number is generated to determine whether a birth or death event transpires. After an individual is added or removed from the population, birth and death rates are updated based on the new population size, and the steps repeat. Extinction occurs at the first time point where the population equaled zero.

Throughout this study, I consider populations that are self limiting. In deterministic models, self-limiting populations experiencing logistic growth reach a stable carrying capacity determined by the intrinsic birth rate (*b*) and the density-dependent death rate (*d*) (Eq. 1).

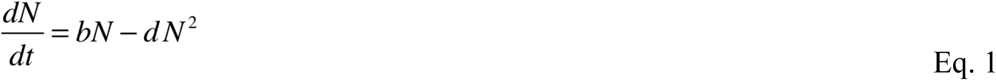

In the stochastic model formulation, births and deaths are modeled as discrete events, also referred to as “reactions” (Anderson and Kurtz 2011). The equivalent birth event rate for logistic growth model is equal to *bN* (Eq. 2):

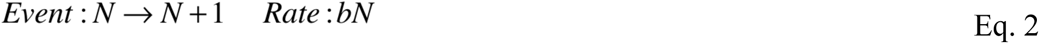

To study effects of mate limitation, I modified the birth event rate to include mate search. Previous work has yielded an equation governing the encounter rate between one individual and other individuals when searching in three-dimensional environments (Gerritsen and Strickler 1977). Results for the two-dimensional case yield qualitatively equivalent results and are shown in the supplementary material. The mate encounter rate is dependent upon the speed at which individuals move (*V*) and the radius at which they can detect a mate (*R*). Here, I assume that males and females move at the same speed and that there is a 1:1 male to female ratio. Multiplying the intrinsic birth rate (*b*) by the probability that at least 1 mate will be encountered yields the following birth event rate for mate-limited populations (Eq. 3):

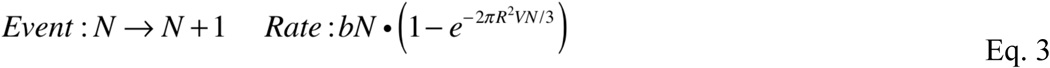

The death rate functions used for the two cases were identical (Eq. 4):

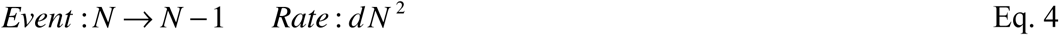

As an illustration of the difference between deterministic and stochastic models, I investigated cases where long-term population dynamics of the various populations would be equivalent in the deterministic case; both mate-limited and non-limited populations would have nearly identical carrying capacities if these dynamics were translated to deterministic models. I simulated population trajectories and evaluated time to extinction using CTMCs. I used the two birth rates expressions above (Eq. 2 and 3) for the scenarios with and without mate limitation, and Eq. 4 for the death rate in all models. I recorded the mean time to extinction (MTE) for 1000 simulated chains (populations).

### Evaluating diversity with island biogeography theory

The theory of island biogeography formalized the concept that long-term community diversity is governed by the rate at which taxa enter the community (i.e. immigration) and the rate that taxa leave the community (i.e. extinction). In island biogeography models, the immigration rate and extinction rate of taxa within a community change as a function of the number of taxa currently present in a community (MacArthur and Wilson 1967). Thus, the expected diversity (defined here as the number of coexisting taxa) of the community is identified by finding the number of taxa where the immigration and extinction rates are equal. To evaluate the accuracy of this analytical approximation for these simulations, I calculated the expected long-term diversity for a community consisting of populations with identical birth and death rates (and, therefore, identical MTEs).

I compared results of the analytical estimates of diversity to simulations of diversity in a stochastic reaction network model (coupled, simultaneous CTMCs) explicitly tracking each population. In the stochastic reaction network, the community-level immigration event rate was a function of current diversity. Immigration events were modeled as a population increasing from 0 to a small population size, in this case of 2 individuals (Eq. 5):

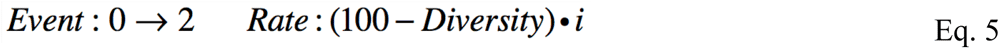

The immigration rate decreased linearly as a function of current community diversity, and reached zero when 100 taxa were present. I used an immigration constant of *i* = 0.001, and conducted these simulations across different parameters governing mate finding.

### Simulation of community structure with heterogeneous taxa

To study how demographic consequences of mate finding scale to communities with heterogeneous populations, I simulated communities where populations had varying intrinsic growth rates (*b*). I evaluated how changes in mate searching parameters affected the diversity, rarity, and dominance of taxa within the simulated communities. Taxa within each community experienced the same degree of mate limitation, which was determined by changing the values of *R* and *V* over the range of 0.5 to 1.2. Across these combinations of search radius and search speed, mate finding is limiting to population growth in small populations, but not limiting in large populations (those with 20 or more individuals).

For each combination of search radius and search speed for the mate limited populations, I simulated a stochastic reaction network where intrinsic birth parameters (*b*) were randomly drawn from a lognormal distribution with a mean of the underlying normal distribution was 0 and the standard deviation was 0.25. The death parameter (*d*) was fixed at 0.1 for all populations. I used a lognormal distribution of growth rates, because a lognormal distribution provides a good fit to the observed abundance distributions of microbial taxa (Shoemaker et al. 2017). Additionally, populations defined by these growth rates routinely become extinct within computationally tractable time scales. However, I verified that simulation results were qualitatively similar when using a normal distribution of birth rates. Again, immigration was a linearly decreasing function of current diversity, where an immigration event was modeled as a change in population size from 0 to 2 (Eq. 5). Immigrant populations were assigned a new birth parameter from the lognormal distribution. After a burn in period of 5 000 000 events, I recorded instantaneous measurements of diversity, dominance (largest population size), and mean population size every 200 000 events. I compared results of simulations where mates were limiting to results of simulations where mates were not limiting.

## Results

### Single population dynamics

First, I compared time to extinction for non-limited populations (i.e. asexual reproducers) and for sexual populations where mates must be encountered. For sexual populations, I evaluated two combinations of search radius and search speed. One scenario indicates a poor searcher with low search radius and speed (*R* and *V* = 0.62), and the other was a more effective searcher with higher search radius and speed (*R* and *V* = 0.8). I chose parameter values that would yield equivalent long-term population dynamics if these populations were modeled deterministically; all three scenarios have nearly identical population densities where the birth rate equals the death rate, indicating equal carrying capacities in the absence of stochasticity (Fig. 1b). The per-capita birth rate is much higher in small populations for the asexually reproducing populations than for sexually reproducing populations (Fig. 1A). However, the birth rate in the sexual populations increased as individuals became more effective at finding mates. Multiplying individual birth rates by population size yields population-level birth minus death rates (Fig. 1B). Effects of mate limitation are prominent at small populations, but negligible as population size increases. With CTMC models, it is also possible to calculate the probability that the next event in the model will be a birth or a death. In models with asexual reproduction, it is highly unlikely that a death will occur in a small population. This probability of population decline at low population sizes is increased when mate limitation is present (Fig. 1C).

**Figure 1:**
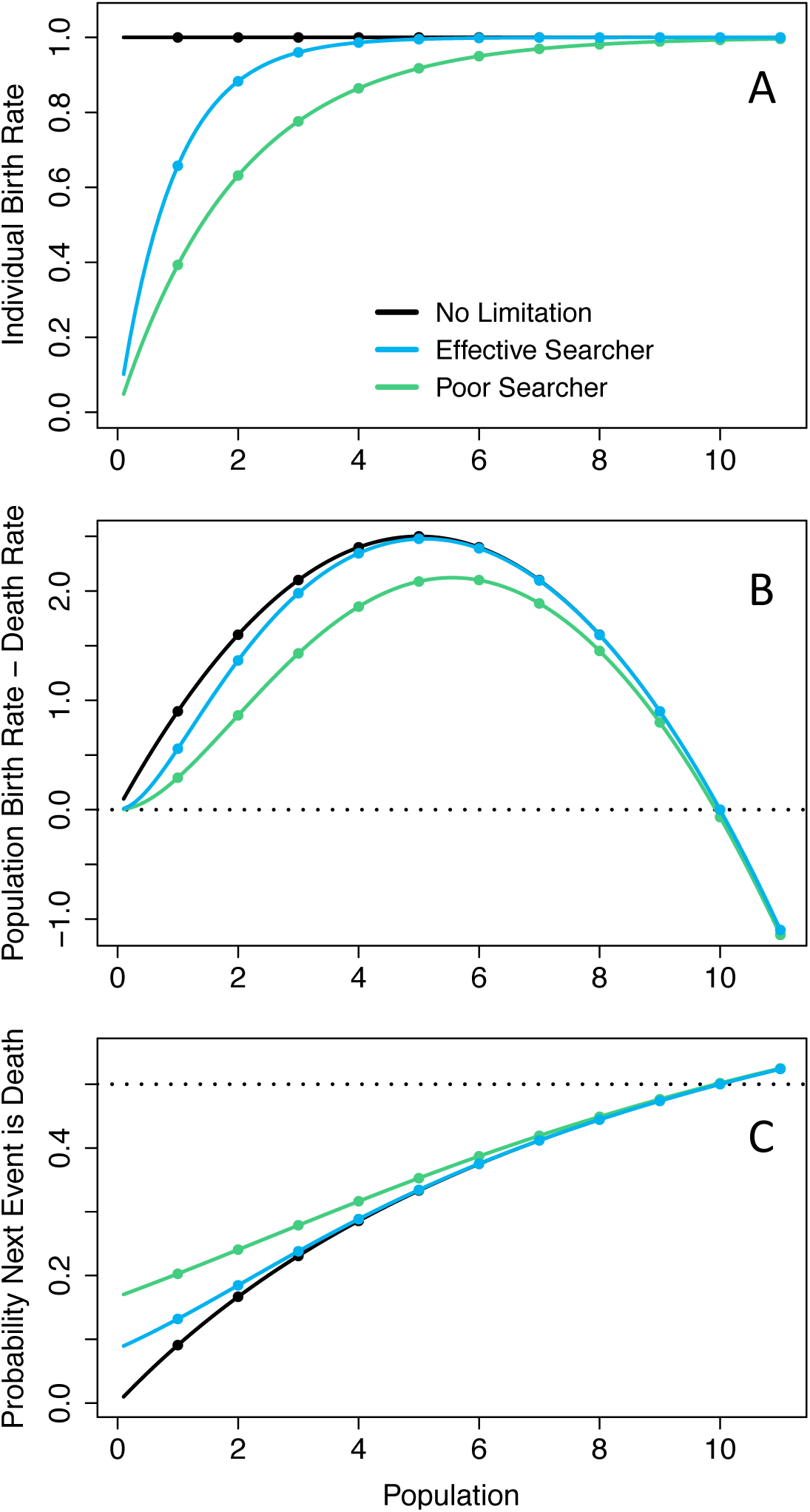
Mate limitation decreases the individual level birth rate at low population density (panel A), which influences both the population-level growth rate (panel B) and the probability that the next event in the model will be a death (panel C). Effects of mate limitation on population growth become negligible as population sizes increase, as visible by the convergence of the three scenarios at larger populations. The effect of mate limitation is the difference between the aseuxally reproducing populations (black line) and the sexually reproducing populations (blue and green lines). Population growth rates are suppressed more strongly in poor mate searchers (green line) than effective mate searchers (blue line). The dashed line in C indicates a probability of 0.5, where a birth and death are equally likely.

I recorded the time to extinction for 1000 simulated populations parameterized with the three birth rate scenarios shown in Fig 1. All populations had equivalent death rates. To evaluate the effect of initial conditions, I used initial population sizes of 10 and 2. Asexual populations persisted longest of the three scenarios, with a MTE of 2792 for an initial population size of 10 and 2664 for an initial population size of 2. In mate-limited populations, MTEs for effective searchers were 1579 and 1557 (for initial size of 10 or 2 individuals), while for poor searchers MTEs were 515 and 477.

Across all populations, rapid extinction (very short MTE) was more common than when the initial population size was 2, rather than 10. The decrease in MTEs between populations with an initial population of 10 and an initial population of 2 is partially due to this higher frequency of very short times to extinction (Fig. 2).

**Figure 2:**
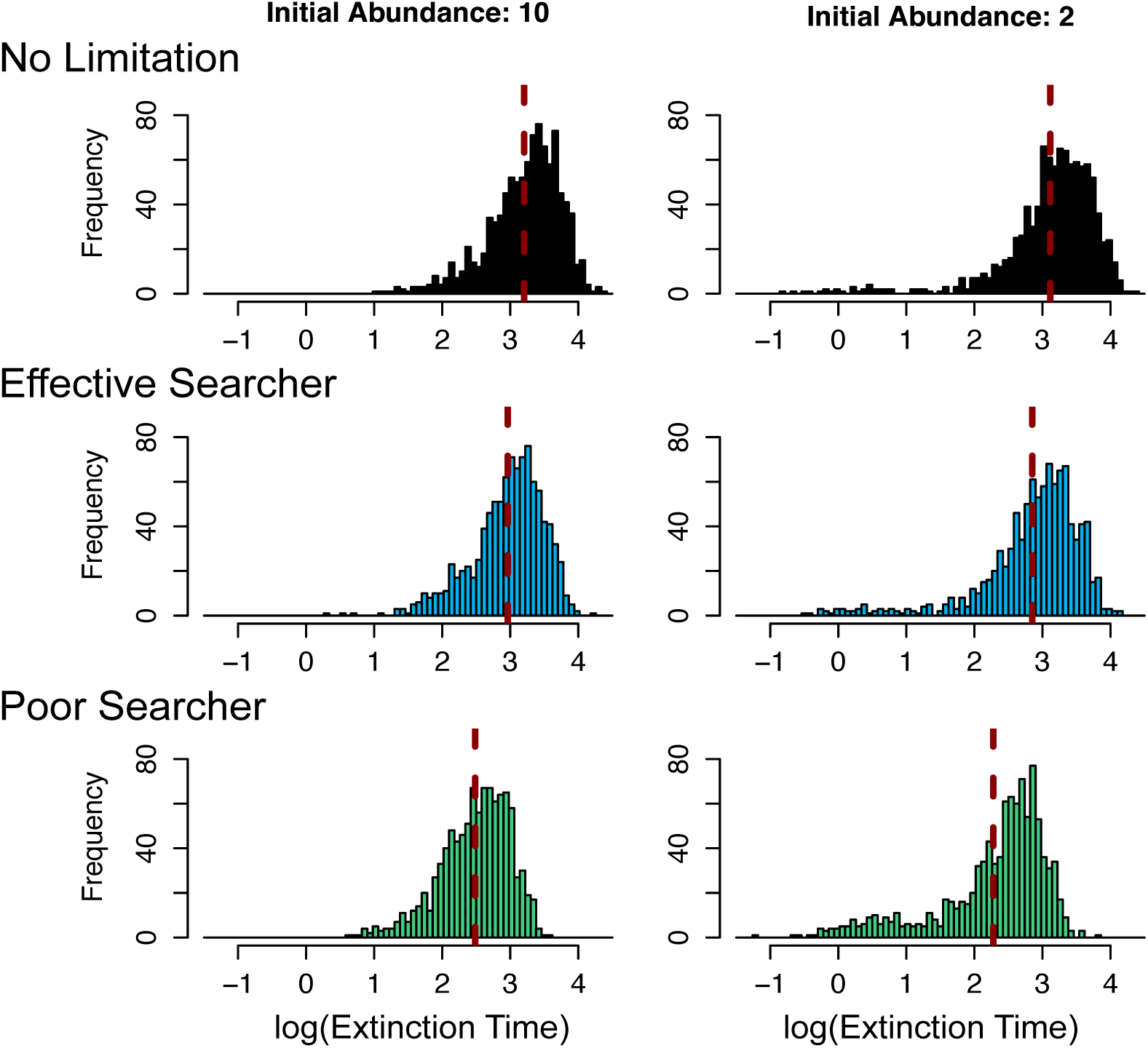
Times to extinction from 1000 simulated populations for communities with no mate limitation (top), mate limitation with effective searchers (midde), and mate limitation with poor searchers (bottom). Mean time to extinction (dashed red lines) decreases as the probability of encountering a mate decreases, and is therefore lowest for poor mate searchers. Time to extintion is also affected by initial population size, and decreases when the population is initiated with 2 individuals (right column) as compared to 10 individuals (left column).

### Evaluating diversity with island biogeography theory

Assuming that a community consisted of populations with identical birth and death rates, I calculated the estimated long-term diversity for the three birth rate scenarios from the associated extinction rates (shown in Fig. 2). The extinction rate for a single population is 1/MTE, meaning the extinction rate for a community of *m* taxa is *m*/MTE. I used the same rate of immigration in each scenario. The immigration rate was a linearly decreasing function of current diversity, and reached 0 when 100 taxa were present (Fig. 3). Thus, no more than 100 taxa could exist in the community. An approximation of long-term diversity under these assumptions can be found using the formula (Eq. 6). For the expected diversity calculations and associated simulations, I used an immigration constant *i* = 0.001, which determines the slope and intercept of the immigration function. However, the stochastic nature of the simulations means that these calculations will be inexact, because the populations never reach equilibrium

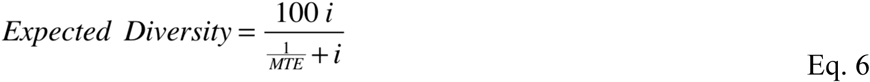

**Figure 3:**
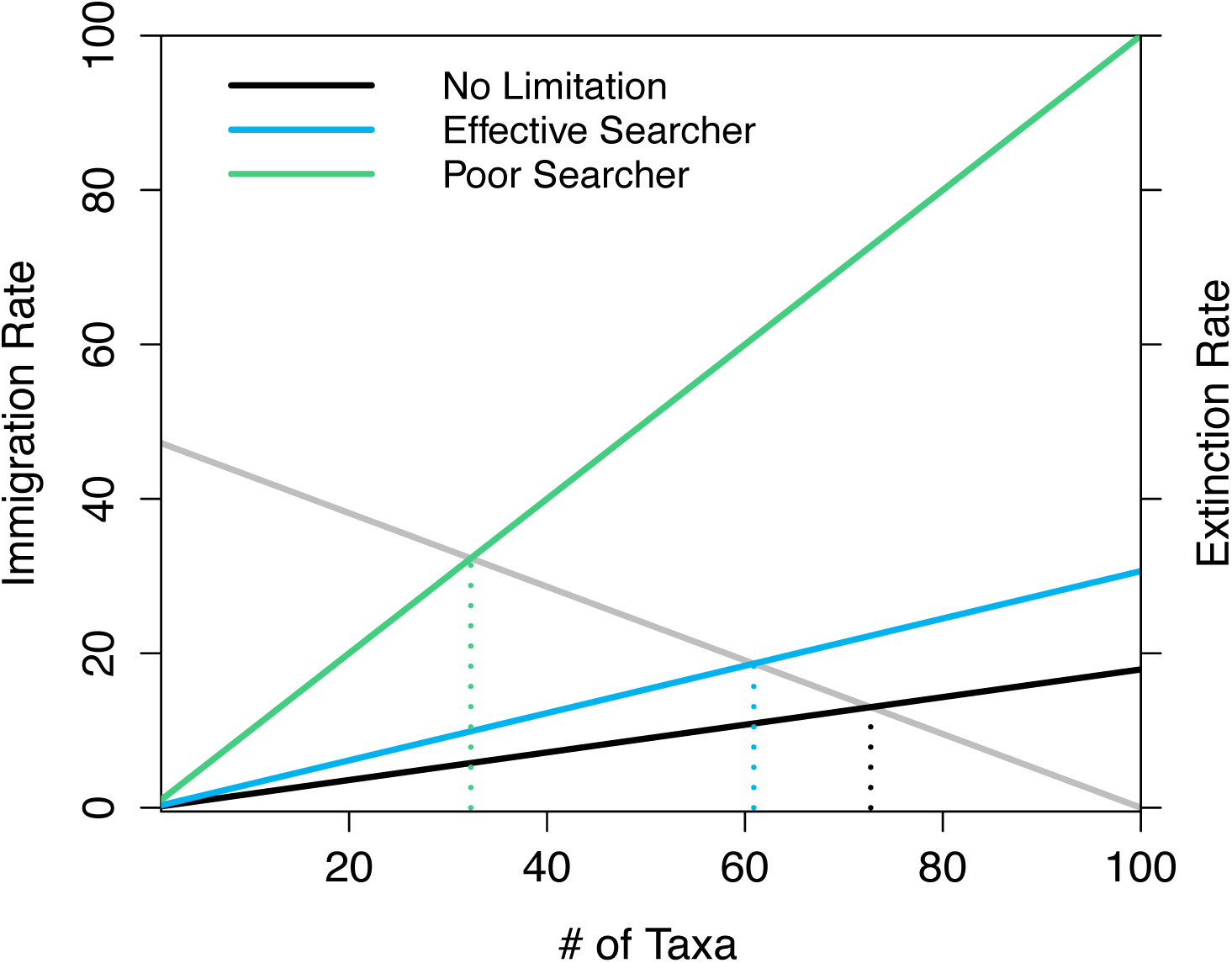
When using the same immigration function (grey line, left axis), mate limitation affects expected diversity by changing time to population extinction. A decreased time to extinction results in a greater slope in the community extinction rate (black, blue, and green lines on right axis). Expected diversity can be found by calculating the number of taxa where the immigration rate and extinction rate intersect (indicated by dashed lines). Communities are most diverse when there is no mate limitation (black lines). When populations are mate-limited, but individuals are effective at finding mates, there is a small decrease in expected diversity (blue lines). When individuals are poor searchers, there is a dramatic decline in diversity due to more rapid extinction (green lines). The time scale at which the immigration and extinction rates are shown here is the MTE of the shortest-lived populations (the poor searcher populations). Using a different time scale alters the y-axes, but does not change where lines intersect.

Eq. 6 shows that the long-term diversity is a function of mean time to extinction. As MTE approaches infinity, the expected diversity approaches the diversity level where immigration is zero (in this case, 100). Conversely, as MTE approaches zero, expected diversity also approaches zero. I evaluated the accuracy of this approximation using explicit simulations of simultaneously coexisting populations using the same parameters. The two estimates of diversity were within one unit (taxon). Approximations using Eq. 6 yielded expected long-term diversities of 72.6, 60.9, and 32.3 for the scenarios of no limitation, effective searching, and poor searching, while simulations yielded long-term average diversities of 73.1, 61.8, and 32.9.

### Simulation of community structure with heterogeneous taxa

Next, I simulated communities in which taxa had varied growth rates. I compared diversity, mean population size, and dominance (population size of most abundant taxon) of communities containing mate-limited taxa and those with non-limited taxa. When sexually reproducing taxa were highly effective searchers due to high search radius and/or speed, mate limitation had little effect on the effective birth rates of those taxa. Then, diversity and population size converged to results from communities containing asexual populations (Fig. 4). However, the abundance of the most dominant population was not affected by search efficacy or reproductive method (Fig. 4). Cells in the figure panels are scaled by the values from the no-limitation simulations (a value of 1 indicates equivalent results).

**Figure 4:**
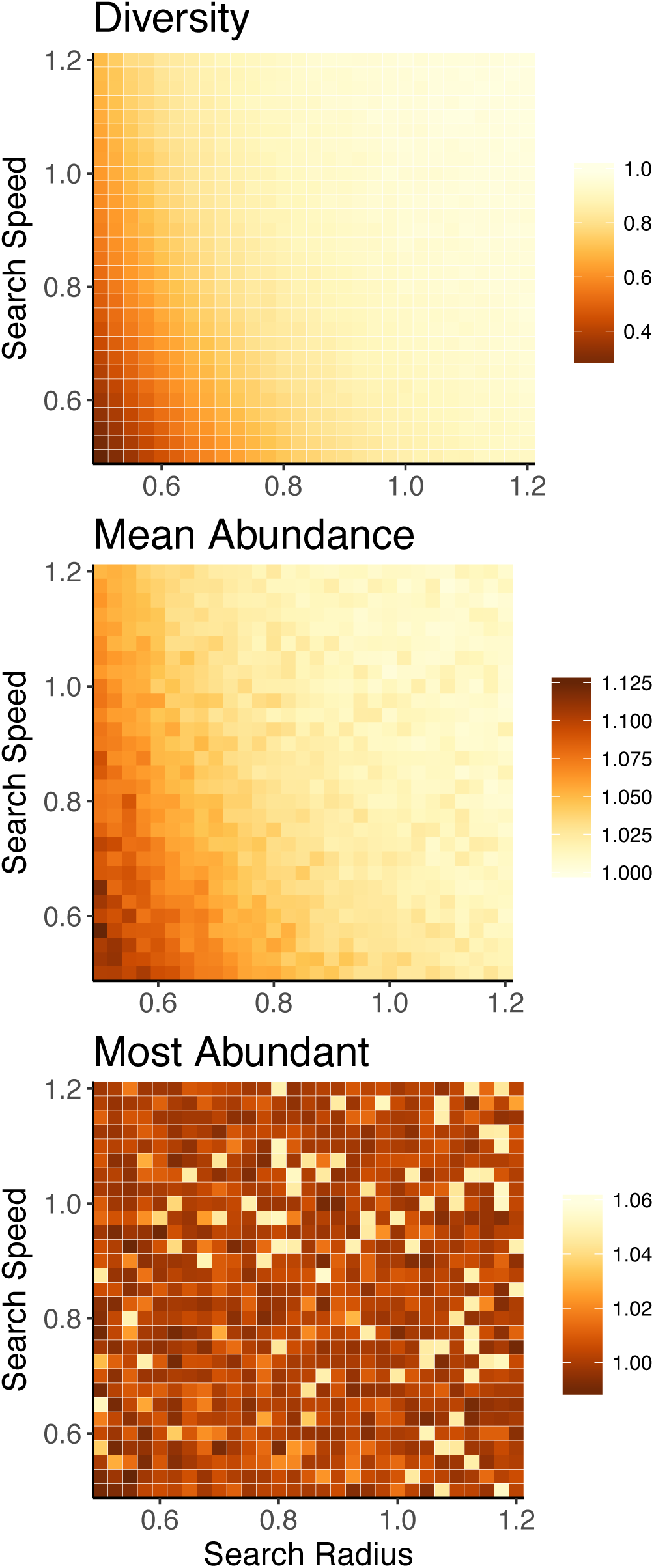
Heatmaps show average diversity (top), abundance (middle), and dominant population size (bottom) from simulations of mate-limited communities where populations have varying search radius and search speed. Cells within each heatmap show the results for communities consisting of populations with the given search radius and search speed. Cell values are scaled by results from communities without mate limitation. Thus, a value of 1 idicates results equivalent to those of non-limited communities. The diversity (top) and mean abundance size (middle) within communities containing mate-limited populations change in response to mate limitation, as stronger mate limitation corresponds to decreased diversity and larger average population size. When the search radius and search speed are large, the probability of finding a mate approaches 1, and results from mate-limited and non-limited communities converge. However, the size of the most abundant population (bottom) is not affected by mate limitation.

Communities containing the poorest mate searchers experienced the greatest declines in diversity, in comparison to the communities with asexual populations. The communities with the strongest mate search limitation (R = 0.5 and V = 0.5) had mean diversity of 17.7 taxa; conversely, communities where mates were not limiting had mean diversity of 68.5 taxa. Similarly, mean population size was 11.2 for the most mate-limited communities, but 10.2 in non-limited communities. Another measurement of rareness, the skewness of population abundances, showed a similar result (see supplementary material). Higher skewness indicates more rare taxa. Average skewness in the distribution of population sizes was 0.18 in communities with greatest mate limitation, and 0.57 in non-limited communities. However, the abundance of the largest population was not consistently related to mate limitation. The abundance of the dominant population in communities with mate-limited populations could be higher or lower than the dominant population in the non-limited community.

Multiple regression analyses showed that mate search speed and search radius explained approximately 90% of variation in diversity and mean abundance in the communities with mate-limited sexual populations (see supplementary material). In contrast, search radius and search speed explain only 1% of variation in dominance (maximum population size). In mate-searching populations, the same degree of limitation could be generated with different combinations of search radius and search speed. Any combination of *R* and *V* that produces a constant value of *VR*^2^ yields an equivalent probability of encountering a mate (Eq. 3).

## Discussion

This study shows that the constraint of mate finding influences emergent community properties, including diversity and average population size. Mate limitation strongly suppresses birth rates when populations are small (Fig. 1), leading to a higher probability that sexually reproducing populations will decline when rare. These discrepancies in birth rate lead to shorter times to extinction in taxa that must find a mate, versus those that reproduce asexually (Fig. 2). This effect is particularly strong when populations are introduced at low density, which is a plausible scenario when considering newly established populations. In stochastic simulations, communities consisting of asexual taxa maintained greater diversity due to a longer expected persistence time of each population (Fig. 3). In the case where immigration is a linear function of current diversity, expected diversity increases as MTE increases (Eq. 6). When these simulations were extended to communities with heterogeneous taxa, differences in diversity were amplified, because mate limitation had especially strong negative effects on taxa with already-low growth rates (Fig. 4). Thus, mate limitation decreased the number of coexisting taxa, primarily by excluding low abundance taxa. However, mate limitation has minimal consequences in larger populations, and therefore the population size of the most abundant taxon was not related to reproductive method or mate search efficacy.

The degree of mate limitation is a function of search ability, which is determined by search radius and search speed. As either search variable (radius or speed) increases, the probability of finding a mate approaches 1, indicating no limitation to the population birth rate. In this case, simulation results of sexual populations with mate finding converge to those of populations without limitation. This is also evident when looking at per capita and population growth curves (Fig. 1). Birth rates asymptotically reach the no-limitation case as mate search becomes more effective. Simulation results are qualitatively similar if instead considering a two-dimensional search (see supplemental material). As in the three-dimensional scenario, mate limitation led to lower diversity and rarity, but had no effect on maximum population size. When searching for mates in three dimensions, the search radius has greater consequence of finding a mate than the search speed. This observation suggests that a trait affecting search radius (such as eyesight) might have a greater fitness effect in a 3D environment than a trait affecting search speed (such as swimming velocity); an incremental increase in search radius would lead to a greater effective birth rate than an increase in search speed. Finally, this study highlights the utility of stochastic models for studying community structure. Deterministic models show equivalent long-term dynamics for populations with the same carrying capacity, whereas these stochastic models show pronounced differences. Thus, this study is concordant with prior models showing that Allee effects increase extinction rates (Brassil 2001, Leibhold and Bascompte 2003, Dennis et al. 2016), and further demonstrates that these population-level effects alter emergent community properties.

Results from these models mirror empirical findings that microbial populations (with asexual reproduction) tend to be high in diversity and rarity, although not distinct from other communities in the dominance of abundant taxa (Locey and Lennon 2016). In these simulated communities, eliminating the constraint of mate finding translated to greater diversity with a higher frequency of low-abundance taxa, while the population size of the most abundant taxon was unaffected (Fig. 4). Thus, allowing for mate finding generates a parsimonious explanation for the community-level patterns observed in comparisons of micro and macro ecological communities. Using empirical data to probe the hypothesis that mate limitation constrains diversity and rarity illustrates the plausibility of this explanation. For example, one study used very deep 16S amplicon sequencing to evaluate whether marine bacterial populations that appeared to only be present seasonally were, instead, consistently present at abundances below the usual detection limit (Caporaso et al. 2012). At a depth of approximately 11 million sequences, 48% of sequences appeared only once (Caporaso et al. 2012). Similarly, another study used deep sequencing of human gut samples to generate rarefaction curves illustrating how many taxa were observed in response to sequencing depth. New taxa continued to be identified after one million sequences were recovered (Turnbaugh et al. 2010). Even if many of the observed rare taxa are the product of sequencing errors, these findings suggest persistence of extremely low abundance taxa (fewer than 1 individual per million). For sexually reproducing populations, these relative abundances could be prohibitively low for individuals to find mates within a lifetime. Finally, asexual reproduction in microbes makes it possible for single individuals to establish populations in new environments. Given the demonstrated plausibility of immigration from microbial seed banks (Lennon and Jones 2011, Caporaso et al. 2012), the growth rate of small populations is especially relevant for the persistence of microbial taxa.

Diversity is a common outcome variable in ecological studies, though there is ongoing debate about how diversity is related to community function (Shade 2017). In the models studied here, diversity is a byproduct of population demographics, including birth rate and mate search ability. More generally, these models show that diversity can be affected by neutral and stochastic processes. Subsequent empirical studies using diversity as an outcome variable might also collect information about immigration and extinction rates to determine whether diversity reflects these processes. For example, surveys of human-associated microbial communities have found variation in diversity across body sites (Caporaso et al. 2011, Huttenhower et al. 2012). Gut and oral bacterial communities are especially diverse (Huttenhower at el. 2012), but these habitats could conceivably have higher immigration rates than other body sites due to daily introduction of bacteria within food (David et al. 2014). Similarly, a recent study found little evidence that fungi can persist within the healthy human gut, but still identified hundreds of fungal taxa in stool samples (Auchtung et al. 2018). The high diversity of fungi in the human gut, in spite of their inability to colonize this habitat, was attributed to persistent immigration of fungi on ingested foods (Auchtung et al. 2018). These studies, coupled with the modeling results presented here, demonstrate how diversity could change independently of community function.

Mathematical models serve as an unbiased line of inference for linking mechanisms to emergent community properties. In the context of comparing community structures of microbial communities and macro-organisms, the differing empirical methods used to study communities at these two scales can generate spurious patterns. Thus, it is hard to discern which findings are true distinctions between micro and macro communities, and which are artifacts of methodology. For instance, DNA sequence similarity is often used to define microbial taxa, whereas macro organisms are generally identified using direct observations. The differences in error rate and detection limits between these two methods could also explain the higher diversity and rarity in microbial communities. Several steps in the workflow of generating 16S amplicon data, including variation in sample processing and sequencing errors, can generate observations of artifactual rare taxa (Fouhy et al. 2016). Furthermore, macro-organisms are often identified using morphological characteristics, but many more taxa can be differentiated if instead using DNA sequencing methods (Fontaneto et al. 2009). Thus, if methodology is a confounding factor when comparing communities, there is uncertainty about whether observed differences are spurious due to sampling bias. Theoretical studies can therefore reinforce empirical findings by determining, with unbiased methodology, whether an identified mechanism can reproduce observed community structure. These modeling results indicate that there are expected differences in diversity and average population size when comparing communities consisting of taxa that reproduce sexually versus asexually.

## Acknowledgements

This work was funded by a postdoctoral fellowship from the Harvard Data Science Initiative. I received valuable feedback from Michael Baym, Anurag Limdi, and members of the Baym lab. The research computing staff at Harvard Medical School provided technical support for model simulations.

## References

Allen LJS, Allen EJ. A comparison of three different stochastic population models with regard to persistence time. Theoretical Population Biology 2003; 64: 439–449.

Allen LJS, Fagan JF, Högnäs G, Fagerholm H. Population extinction in discrete-time stochastic population models with an Allee effect. Journal of Difference Equations and Applications 2005, 11, 273–293.

Anderson DF, Kurtz TG. Continuous Time Markov Chain Models for Chemical Reaction Networks. Design and Analysis of Biomolecular Circuits. 2011. Springer, New York, NY, pp 3–42.

Assaf M, Meerson B. Extinction of metastable stochastic populations. Physical Reviews E 2010; 81: 021116.

Auchtung TA, Fofanova TY, Stewart CJ, Nash AK, Wong MC, Gesell JR, et al. Investigating Colonization of the Healthy Adult Gastrointestinal Tract by Fungi. mSphere 2018; 3: e00092–18.

Boukal DS, Berec L. Single-species Models of the Allee Effect: Extinction Boundaries, Sex Ratios and Mate Encounters. Journal of Theoretical Biology 2002; 218: 375–394.

Brassil CE. Mean time to extinction of a metapopulation with an Allee effect. Ecological Modelling 2001; 143: 9–16.

Caporaso JG, Lauber CL, Costello EK, Berg-Lyons D, Gonzalez A, Stombaugh J, et al. Moving pictures of the human microbiome. Genome Biology 2011; 12: R50.

Caporaso JG, Paszkiewicz K, Field D, Knight R, Gilbert JA. The Western English Channel contains a persistent microbial seed bank. The ISME Journal 2012; 6: 1089–1093.

Consortium THMP, Huttenhower C, Gevers D, Knight R, Abubucker S, Badger JH, et al. Structure, function and diversity of the healthy human microbiome. Nature 2012; 486: 207–214.

Courchamp F, Clutton-Brock T, Grenfell B. Inverse density dependence and the Allee effect. Trends in Ecology & Evolution 1999; 14: 405–410.

David LA, Maurice CF, Carmody RN, Gootenberg DB, Button JE, Wolfe BE, et al. Diet rapidly and reproducibly alters the human gut microbiome. Nature 2014; 505: 559–563.

Dennis B. Allee effects: population growth, critical density, and the chance of extinction. Natural Resource Modeling 1989; 3: 481–538.

Dennis B. Allee Effects in Stochastic Populations. Oikos 2002; 96: 389–401.

Dennis B, Assas L, Elaydi S, Kwessi E, Livadiotis G. Allee effects and resilience in stochastic populations. Theoretical Ecol 2016; 9: 323–335.

Dobramysl U, Mobilia M, Pleimling M, Täuber UC. Stochastic population dynamics in spatially extended predator–prey systems. Journal of Physics A: Mathematical and Theoretical 2018; 51: 063001.

Fontaneto D, Kaya M, Herniou EA, Barraclough TG. Extreme levels of hidden diversity in microscopic animals (Rotifera) revealed by DNA taxonomy. Molecular Phylogenetics and Evolution 2009; 53: 182–189.

Fouhy F, Clooney AG, Stanton C, Claesson MJ, Cotter PD. 16S rRNA gene sequencing of mock microbial populations-impact of DNA extraction method, primer choice and sequencing platform. BMC Microbiology 2016; 16: 123.

Gascoigne J, Berec L, Gregory S, Courchamp F. Dangerously few liaisons: a review of mate-finding Allee effects. Population Ecology 2009; 51: 355–372.

Gerritsen J. Sex and Parthenogenesis in Sparse Populations. The American Naturalist 1980; 115: 718–742.

Gerritsen J, Strickler JR. Encounter Probabilities and Community Structure in Zooplankton: a Mathematical Model. Journal of the Fisheries Research Board of Canada 1977; 34: 73–82.

Gillespie DT. Exact stochastic simulation of coupled chemical reactions. The journal of physical chemistry, 1977; 81: 2340–2361.

Gillespie DT. Stochastic Simulation of Chemical Kinetics. Annual Review of Physical Chemistry 2007; 58: 35–55.

Hansen GJA, Carey CC. Fish and Phytoplankton Exhibit Contrasting Temporal Species Abundance Patterns in a Dynamic North Temperate Lake. PLOS ONE 2015; 10: e0115414.

Hutchinson GE. The paradox of the plankton. The American Naturalist 1961; 882: 137–145.

Lande R. Risks of Population Extinction from Demographic and Environmental Stochasticity and Random Catastrophes. The American Naturalist 1993; 142: 911–927.

Lennon JT, Jones SE. Microbial seed banks: the ecological and evolutionary implications of dormancy. Nature Reviews Microbiology 2011; 9: 119–130.

Ley RE, Peterson DA, Gordon JI. Ecological and Evolutionary Forces Shaping Microbial Diversity in the Human Intestine. Cell 2006; 124: 837–848.

Liebhold Andrew, Bascompte Jordi. The Allee effect, stochastic dynamics and the eradication of alien species. Ecology Letters 2003; 6: 133–140.

Locey KJ, Lennon JT. Scaling laws predict global microbial diversity. PNAS 2016; 113: 5970–5975.

Lynch MDJ, Neufeld JD. Ecology and exploration of the rare biosphere. Nature Reviews Microbiology 2015; 13: 217–229.

MacArthur RH, Wilson EO. The Theory of Island Biogeography. 1967. Princeton University Press.

Meyer KM, Memiaghe H, Korte L, Kenfack F, Alonso A, Bohannan BJM. Why do microbes exhibit weak biogeographic patterns? The ISME Journal 2018.

Nemergut DR, Schmidt SK, Fukami T, O’Neill SP, Bilinski TM, Stanish LF, et al. Patterns and Processes of Microbial Community Assembly. Microbiology and Molecular Biology Reviews 2013; 77: 342–356.

Odum HT, Allee WC. A Note on the Stable Point of Populations Showing Both Intraspecific Cooperation and Disoperation. Ecology 1954; 35: 95–97.

Purvis A, Gittleman JL, Cowlishaw G, Mace GM. Predicting extinction risk in declining species. Proceedings of the Royal Society of London B: Biological Sciences 2000; 267: 1947–1952.

Shoemaker WR, Locey KJ, Lennon JT. A macroecological theory of microbial biodiversity. Nature Ecology & Evolution 2017; 1: 0107.

Shade A. Diversity is the question, not the answer. The ISME Journal 2017; 11: 1–6.

Stephens PA, Sutherland WJ, Freckleton RP. What Is the Allee Effect? Oikos 1999; 87: 185–190.

Stephan T, Wissel C. Stochastic extinction models discrete in time. Ecological Modelling 1994; 75–76: 183–192.

Sun G-Q. Mathematical modeling of population dynamics with Allee effect. Nonlinear Dynamics 2016; 85: 1–12.

Turnbaugh PJ, Quince C, Faith JJ, McHardy AC, Yatsunenko T, Niazi F, et al. Organismal, genetic, and transcriptional variation in the deeply sequenced gut microbiomes of identical twins. PNAS 2010; 107: 7503–7508.

Volterra V. Population growth, equilibria, and extinction under specified breeding conditions: a development and extension of the theory of the logistic curve. Human Biology 1938; 10: 1–11.

Xu Z, Knight R. Dietary effects on human gut microbiome diversity. British Journal of Nutrition 2015; 113: S1–S5.

